# Lipids modulate open probability of RyR1 under cryo-EM conditions

**DOI:** 10.1101/2024.12.20.629683

**Authors:** Chenyao Li, Rouslan G. Efremov

## Abstract

Ryanodine receptors (RyRs) are intracellular tetrameric ion channels responsible for Ca^2+^ release from the sarcoplasmic and endoplasmic reticulum. Among the three known mammalian RyR isoforms, RyR1 is critical for muscle contraction and has been studied most extensively. The cytoplasm-exposed multidomain fragment of RyRs integrates multiple cellular signals that regulate channel gating and small deviations from the physiological open probability of RyRs leads to life-threatening diseases. While cryo-EM has been instrumental in revealing near-atomic details of RyR gating mechanisms, the open probability of RyR1 under cryo-EM conditions is notably lower than that observed in electrophysiological studies, complicating structural investigations of RyR1 gating modulation. Here, we present a cryo-EM study examining the open probability of RyR1 solubilized in CHAPS with varying lipid concentrations. We found that increasing lipid concentration from 0.001% to 0.05% raised the RyR1 open probability from 16 to 84%; however, RyR1 reconstituted into lipid nanodiscs remained closed. We modelled 72 lipid molecules in the map reconstructed at the highest lipid concentration. These findings demonstrate critical role of lipids in modulating RyR1 gating under cryo-EM conditions and suggest optimal lipid-mimetics for structural studies of RyR1 gating modulation.

## Introduction

Structural studies of ion channels are often performed on the detergent-solubilized proteins ^1–8^. Alternative systems include lipid nanodiscs, amphipols and since more recently lipid liposomes and native membranes ^9–17^. Typically, structural studies aim at determining the structures of the ion channels in several functional conformations such as closed, open, deactivated or primed states ^5–8,16–18^. However, the physiological function of ion channels also involves fine regulation of the channel open probability which is modulated by interactions with soluble or membrane-embedded molecules (including proteins) or by changes in lipid membrane state including trans-membrane electrochemical potential and lipid composition ^18–22^. Such modulations can be studied using single-particle cryo-EM, which, in addition to resolving structures of functional conformations, provides particle counts for each conformation. From these counts, the relative occupancies of the conformations can be determined. The studies of channel gating based on cryo-EM data aim at establishing contributions from individual factors such as binding of small molecules and proteins, post-translational modifications, temperature, and pH to channel gating. Additionally, gating is modulated by lipid mimetics in a channel-specific manner ^22–26^.

The ryanodine receptors (RyRs) are homo tetrameric Ca^2+^ channels with a molecular weight of around 2.2 MDa ^27–29^. They reside in the sarcoplasmic or endoplasmic reticulum (SR/ER) membrane where they mediate calcium signaling in an isoform-dependent manner ^27–31^. Among three RyR isoforms found in mammals, RyR1 is primarily expressed in skeletal muscles, RyR2 in the heart, and RyR3 is expressed at low levels in various tissues ^27–29^.

RyR1 play a crucial role in excitation-contraction (E-C) coupling, thus Ca^2+^ release from SR to cytoplasm through RyR1 triggers muscle contraction ^32,33^. Hence, precise control of RyR1 gating is essential for normal physiological function whereas Ca^2+^ leakage can lead to diseases such as malignant hyperthermia (MH) and central core disease (CCD) ^34–37^. The gating of RyR1 is regulated by both modulators and the membrane environment. While much research has been conducted on regulatory mechanisms of proteins and small molecules ^8,21,38–43^, fewer studies have explored how membrane mimetics influence RyR1 open probability ^24,44,45^. Because even subtle changes in RyR1 gating leads to disease, understanding how individual factors, including membrane mimetics, finetune the open probability of RyR1 is essential.

Previous studies have observed that open probability of RyR1 derived from analysis of single particle cryo-EM data of detergent-solubilized RyR1 (∼50%) was about twice lower than nearly 100% open probability observed in single-channel recordings under otherwise identical conditions ^8^. Curiously, the open probability of RyR1 reconstituted into liposomes estimated by cryo-EM was still low (37% and 62%) ^44,45^. Our previous study based on H^3^-ryanodine binding assays indicated that a mixture of detergent CHAPS with lipids at various concentrations tunes open probability of RyR1 ^24^. This observation has not been verified using structural data and the origin of such a modulation remains unclear.

To further understand the modulation of open probability of RyR1 by lipid mimetics, we investigated the influence of lipids on RyR1 open probability using cryo-EM. We show that changing amount of lipids added to detergent-solubilized RyR1 modulates open fraction of RyR1 under cryo-EM conditions between 16 and 84%. The reconstructions obtained at high lipid concentration allowed us to build models of 18 lipid molecules per RyR1 protomer and visualize the details of RyR1-lipid interactions. Structural analysis suggests that lipids modulate gating by establishing a hydrophobic barrier between the transmembrane domain and the hydrophilic buffer.

These results indicate that the lipid concentration in the detergent micelles is a critical determinant of the channel open probability and must be thoroughly controlled in the experiments seeking to study modulation of RyR1.

## Results

### RyR1 structures with low and high concentration of lipids

Previous studies of RyR1 by cryo-EM utilized 0.001% of lipids (prior to protein concentrating) in the protein solution solubilized in 0.2-0.25% of CHAPS ^8,45^. Under these conditions, RyR1’s open probability estimated from the number of particles assigned to different conformations, was between 16^45^ and 50%^8^ (see Discussion for possible reasons of the discrepancy). Our previous biochemical studies indicated that open probability of solubilized RyR1 depends on the concentration of lipids ^24^. To understand how amount of lipids in solubilized protein solution changes the structure and open probability of RyR1, we solved cryo-EM structures of the protein purified using RyR-specific nanobody Nb9657 ^45^ and plunge frozen it in buffer containing 0.2% CHAPS and either 0.01% or 0.05% POPC (x10 and x50 fold higher lipid concentrations). The cryo-EM datasets were collected in the presence of 50 μM free Ca^2+^, 2 mM ATP, and 5 mM caffeine — conditions that stabilize the open state of RyR1 with Po of 0.93 in single-channel electrophysiological recordings ^8^.The elevated lipid concentration reduced contrast of RyR1 particles in cryo-EM micrographs, with the 0.05% POPC dataset exhibiting a more pronounced loss of contrast (**Figure 1a, b**). Nonetheless, reconstructions of primed and open states were determined to resolution between 3.3 and 3.6 Å for both datasets (**Figure 1, Supplementary Figs 1-2, Supplementary Table 1**) which further referred to as DT-0.05-primed, DT-0.05-open, DT-0.01-primed, and DT-0.01-open. Here DT indicates that the protein was solubilized in detergent. Bound Nb9657 and endogenous co-purified FKBP12 were well-resolved in the density maps (**Figure 1e**). Both datasets were processed using the same protocol and the open probability estimated from the number of particles assigned to primed and open states was 67 and 84% for DT-0.01 and DT-0.05, respectively.

**Figure 1.**
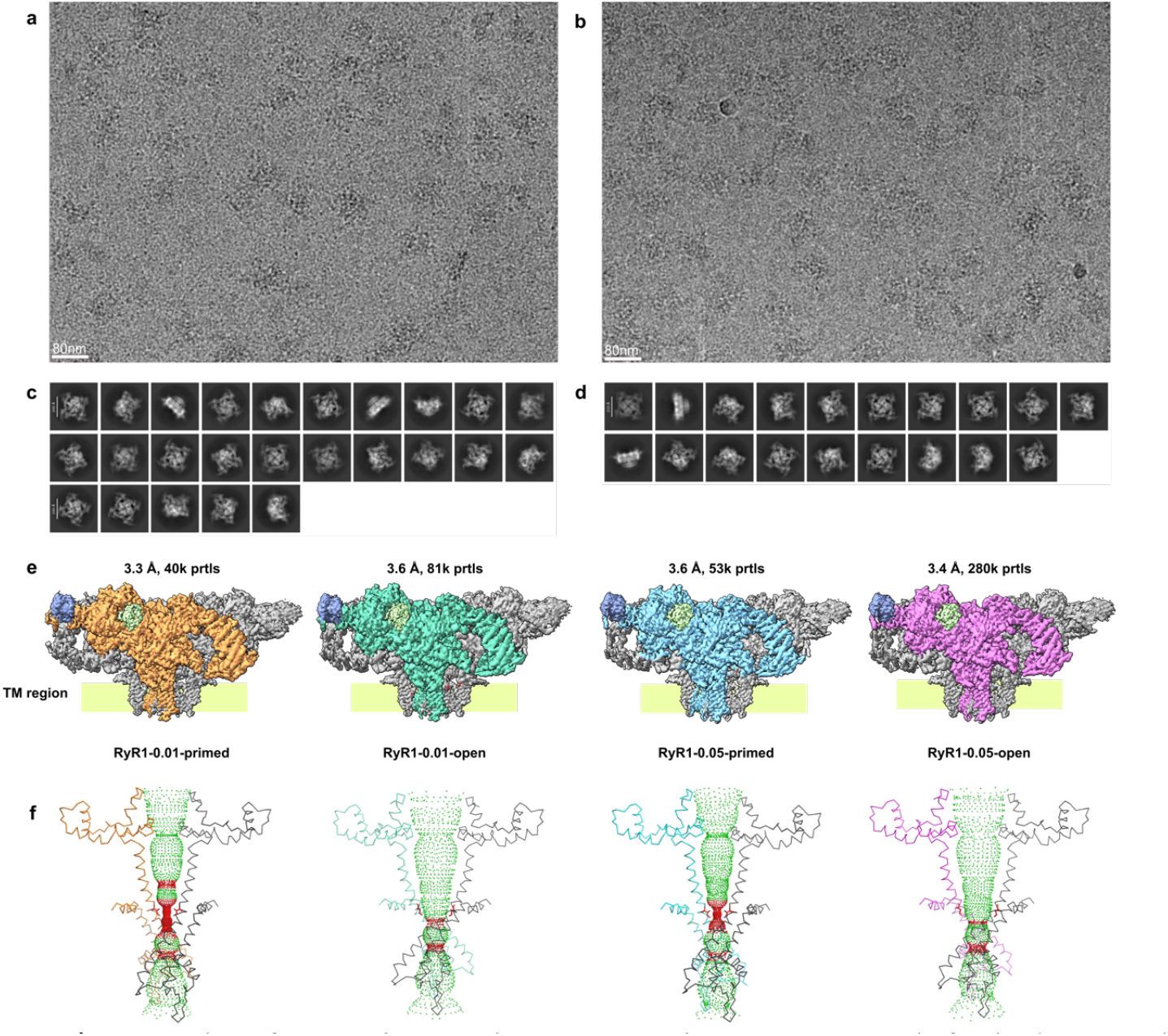
Cryo-EM data of RyR1 with 0.01 and 0.05% POPC. **a, b** Cryo-EM micrographs for the datasets with 0.01% and 0.05% POPC, respectively. **c, d**, Corresponding 2D class averages for each dataset. **e**, 3D density maps of the structures determined from the two datasets, from left to right: DT-0.01-primed, DT-0.01-open, DT-0.05-primed, and DT-0.05-open. The nanobody is shown in blue, FKBP12 in light green, and one monomer from each structure is colored orange, spring green, cyan and magenta, respectively. **f**, Side view of the pore region for each structure showing two protomers—one in grey and the other colored as in **e**.The gate residue, I4937, is highlighted in red. The solvent-accessible surface of the channel, calculated using HOLE ^46^, is shown in green for channel regions with a radius above 4 Å and in red for those below 4 Å.

### The fraction of open RyR1 particles is modulated by lipids

Two new datasets complement previously reported structures determined with 0.001% POPC (DT-0.001 data set), reconstituted into lipid nanodiscs (ND data set) and in POPC liposomes (LP data set). These 5 datasets were collected with RyR1 purified following similar protocol and using the same detergent and lipids, the channel was stabilized in open state using the same channel activators and data were processed using the same protocols as for the two datasets described above these data allow for comparison of open channel probability dependence on lipid mimetics.

We observe that lipid mimetics influence the fraction of particles in open channel conformation. The open probability of RyR1 solubilized in CHAPS is strongly modulated by lipids and increases from 16 to 67 and to 84% as lipid concentration grows from 0.001 to 0.01 and to 0.05% POPC, respectively (**Figure 2, Supplementary Table 2**). Somewhat counter-intuitive observation was that in the lipid mimetics that is considered to better approximate lipid bilayer such as lipids nanodiscs and liposomes, open probabilitiy, Po, did not match values determined by electrophysiology. Thus, RyR1 reconstituted into the lipid liposomes is expected to be nearly fully open (Po >90%) with activators used in the cryo-EM studies (Ca^2+^, ATP, and caffeine) ^47,48^. Thus, for RyR1 reconstituted into lipid nanodiscs, 3D classification consistently failed to identify open conformation, suggesting that RyR1 remained fully closed. Open fraction of RyR1 reconstituted into lipid liposomes (62%)^45^ was lower than that of DT-0.01 (67%) and DT-0.05 (84%) (**Figure 2, Supplementary Table 2**).

**Figure 2.**
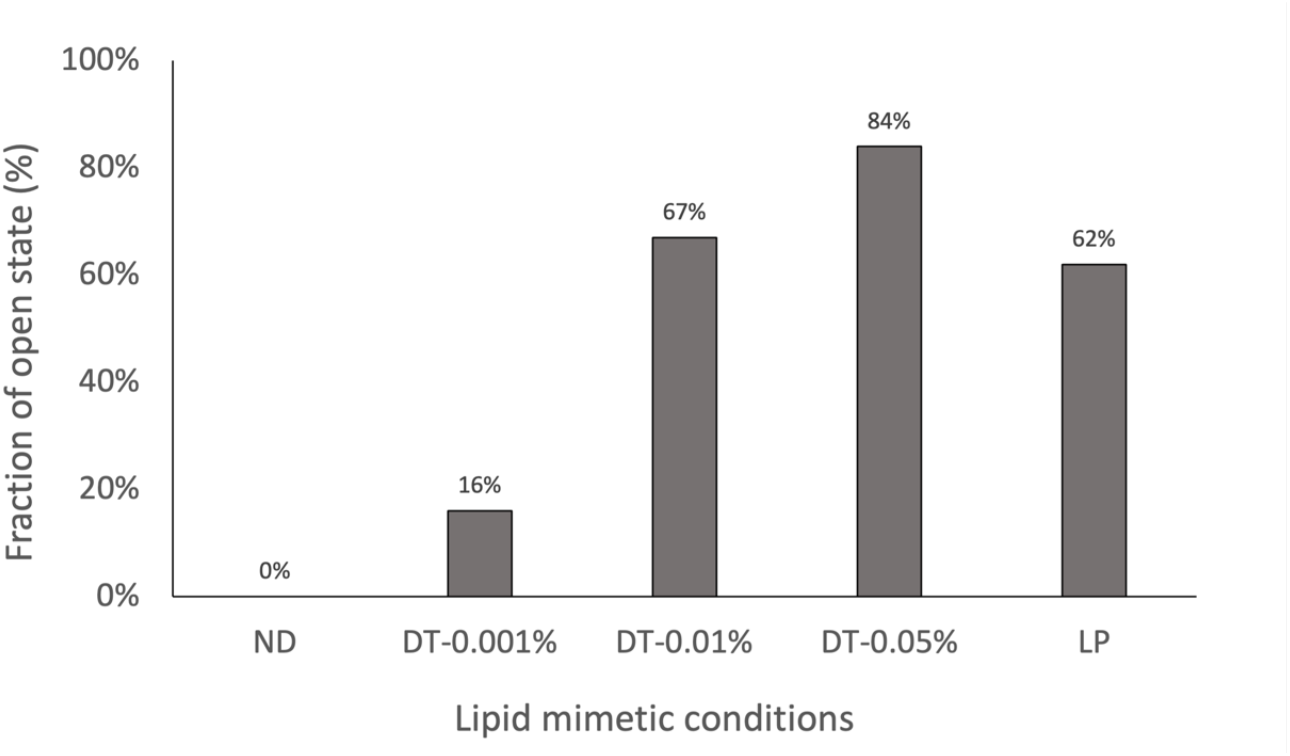
Fraction of open RyR1 on different lipid mimetics. From left to right, ND - RyR1 in nanodiscs, DT – RyR1 in CHAPS detergent micelles supplemented with lipids at indicated concentrations, and in LP - RyR1 reconstituted into POPC liposomes.

### Lipid mimetics influence the channel gate diameter

Structural comparison of primed and open conformations shows that overall conformation of the primed and open states was similar in various lipid mimetic conditions (**Supplementary Figure 3a, b**). However, closer structural analysis revealed that the gate diameters does slightly depend on the lipid mimetics. Thus, in the primed state, the Cα-Cα distance for gate residue I4937 was 11.2 Å for ND, 11.7 Å for all DT, and 12.2 Å for LP models. Different lipid mimetics introduced small but consistent changes in gate diameters for the open states too (Supplementary Figure 3 c, d). The Cα-Cα distances for gate residue I4937 were measured 17.5, 18.5, 19.1, and 18 Å for DT-0.001, DT-0.01, DT-0.05, and LP, respectively. (**Supplementary Figure 3c, d**). These small changes in the gate diameter correlate with variations in the percentage of the particles in open conformation across the investigated lipid mimetics.

### A belt of ordered annular lipids is resolved at high lipid concentration

The map of DT-0.05-open resolved multiple elongated densities into which 72 lipids (18 per protomer named L0-L17) were modelled (**Figure 3**). At the same time, no density for ordered CHAPS molecules which are steroid derivatives with large flat structures composed of four fused rings clearly distinguishable from phospholipids, was observed despite high CHAPS concentration (molar ratios CHAPS: lipids were in the range 250:1 - 5:1). Even though DT-0.05-open map had lower resolution than the maps solved at lower lipid concentrations (DT-0.001-primed or ND-primed), the lipid densities were better defined and had unambiguous connectivity (**Supplementary Figure 4**).

**Figure 3.**
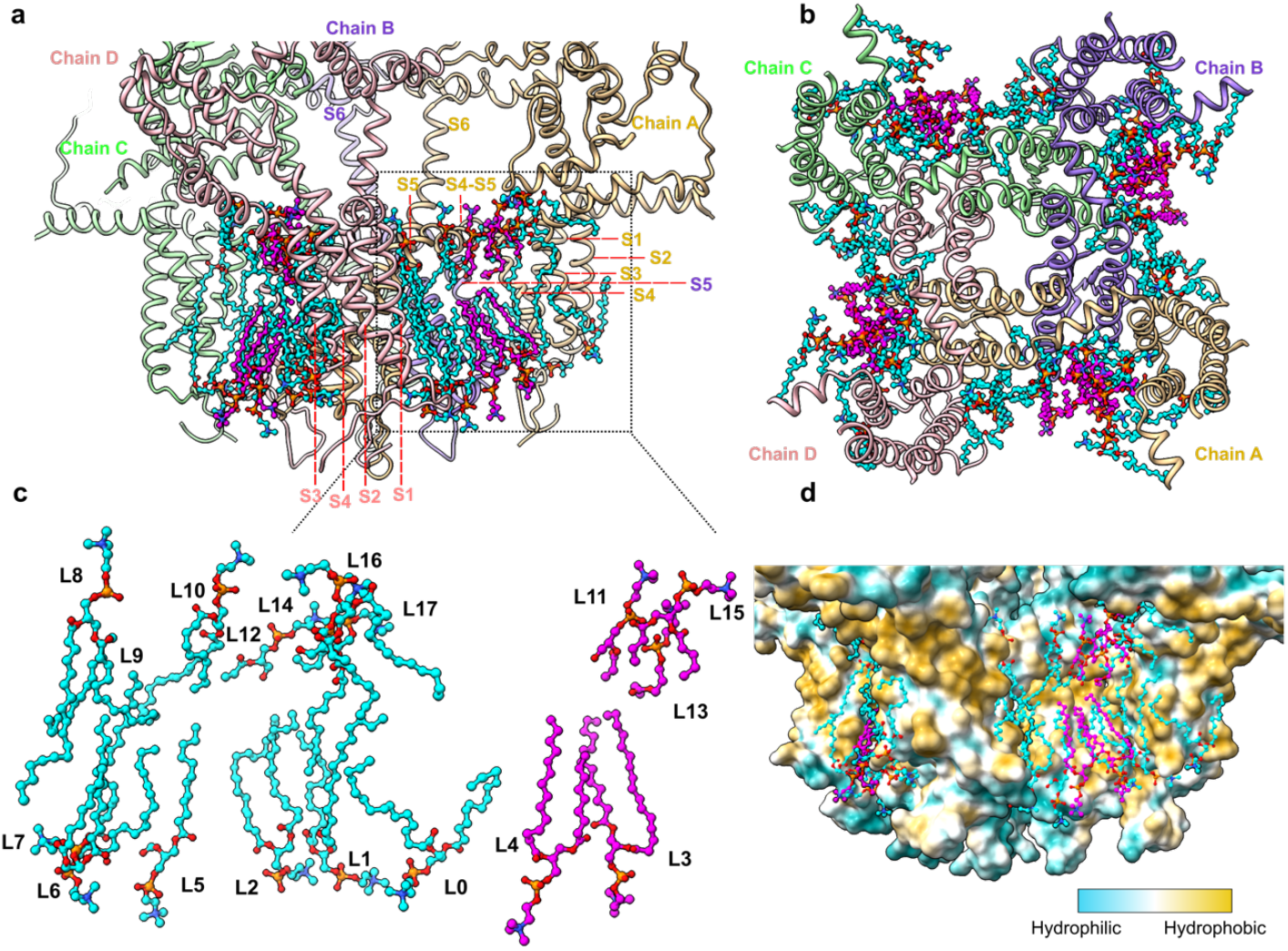
Ordered lipids modeled in the RyR1 structure. **a, b**, Side and top views of POPC modelled into DT-0.05-Open map. The first-layer lipids are colored in cyan, and the second-layer lipids in magenta. The RyR1 structure is presented as a cartoon, and lipids are shown in ball-and-stick representation. **c**, An enlarged view of the modelled lipids. For better visualization, the second layer (right side) is separated from the first (left side). **d**, Lipid positions relative to RyR1 surface colored by hydrophobicity.

The lipids were modeled as POPC, hence it was the only lipid added during the protein preparation. The modeled lipids cover nearly completely the concave regions of trans-membrane domain formed between the 4-helical voltage sensor-like helical bundles (S1-S4) arranged in a cross-like pattern (**Figure 3**). This is an improvement in comparison to the published RyR structures that typically include only two lipids ^49^, while the structural model of closely related IP3R includes seven ^18^.

The modelled lipids are organized in two layers around the surface of membrane embedded RyR1 region. The annular layer of lipids interacts directly with RyR1 (**Supplementary Figure 4**) and accounts for 13 lipids and lipids fragments. Among the lipids with modelled head groups, 3 are located on the luminal leaflet and 9 on the cytoplasmic **(Figure 3 a, c)**. The second layer of lipids contains 5 lipids and lipid fragments and as the 1^st^ layer fills the concave surface of trans-membrane domain with 3 cytoplasmic and 2 luminal lipids (magenta in **Figure 3**).

The details of lipid-RyR1 interactions are shown in **Supplementary Figure 4**. The annular lipids interact extensively with the protein vial hydrophobic interactions and several putative polar contacts with RyR1 amino acid residues located in the interface between hydrophobic and polar surfaces of the trans-membrane domain **(Supplementary Figure 5)**. Lipids from the second layer, on the other hand, interact only with annular lipids except for L3 that forms a putative hydrogen-bond with Y4912 **(Supplementary Figure 4b)**.

To analyze details of lipid interactions with RyR1, we conducted a protein-ligand interaction analysis using the Protein-Ligand Interaction Profiler ^50^. In total, 49 residues per protomer interact with lipids (**Supplementary Figure 4**), all are conserved between rabbit and human RyR1. Among these residues, mutations R4824C (R4825 in humans), A4845V (A4846 in humans), and F4859 deletion (F4860 in humans) (**Supplementary Figure 5**) have been associated with Central Core Disease, whereas the mutation A4845V was linked to Myotonic Dystrophy Type 1 ^51,52^.

To understand the role of lipids in stabilizing open RyR1 conformation, we compared the density of lipids between the maps obtained from detergent solubilized RyR1 prepared with different lipid concentrations and RyR1 reconstituted into nanodiscs. Due to the low resolution, maps from RyR1-LP and DT-0.001-open were not used in the analysis. When compared to DT-0.05-open map, the maps obtained at lower lipid concentrations had lipid densities in the same positions but were much weaker and more fragmented (Supplementary Table 3). The cross-correlation values of the modeled lipids followed a similar trend as the density (Supplementary Table 4). Lipids in the first layer were conserved, except for L5, L12, L8 and L16 while density for the lipids in the second layer was too weak to model lipids reliably. This suggest that increasing lipid concentration leads to populating lipids in the weaker binding sites around the trans-membrane domain making consistently a lipid-bilayer-like structure between lobes formed by S1-S4 helical bundles.

### Structural adaptations of lipid during primed to open conformational transition

The conformational changes in RyR1 trans-membrane domain during transition from closed/primed to open state include rotation of S1-S4 lobes by ∼2 degrees outwards and expansion of the cytoplasmic side of pore-forming S5-S6 helices with amplitude of around 3 Å (**Supplementary Video 1**). To understand the involvement of lipids in this transition and their possible role in stabilizing open state conformation, we compared the RyR1-lipid interactions between the DT-0.05-open state and DT-0.05-primed states. Most of the lipids and protein-lipid interactions are conserved between these states with exception of L4-5 and L11-13, L14-15, which might be less well resolved due to lower resolution (**Figure 4 a**). Only minor structural rearrangements in lipids L2, L3, L7, L16, and L17 were observed. The lipid positions adapt to the altered membrane-exposed surface of the trans-membrane domain without global positional rearrangement (**Figure 4 b, Supplementary Video 1**).

**Figure 4.**
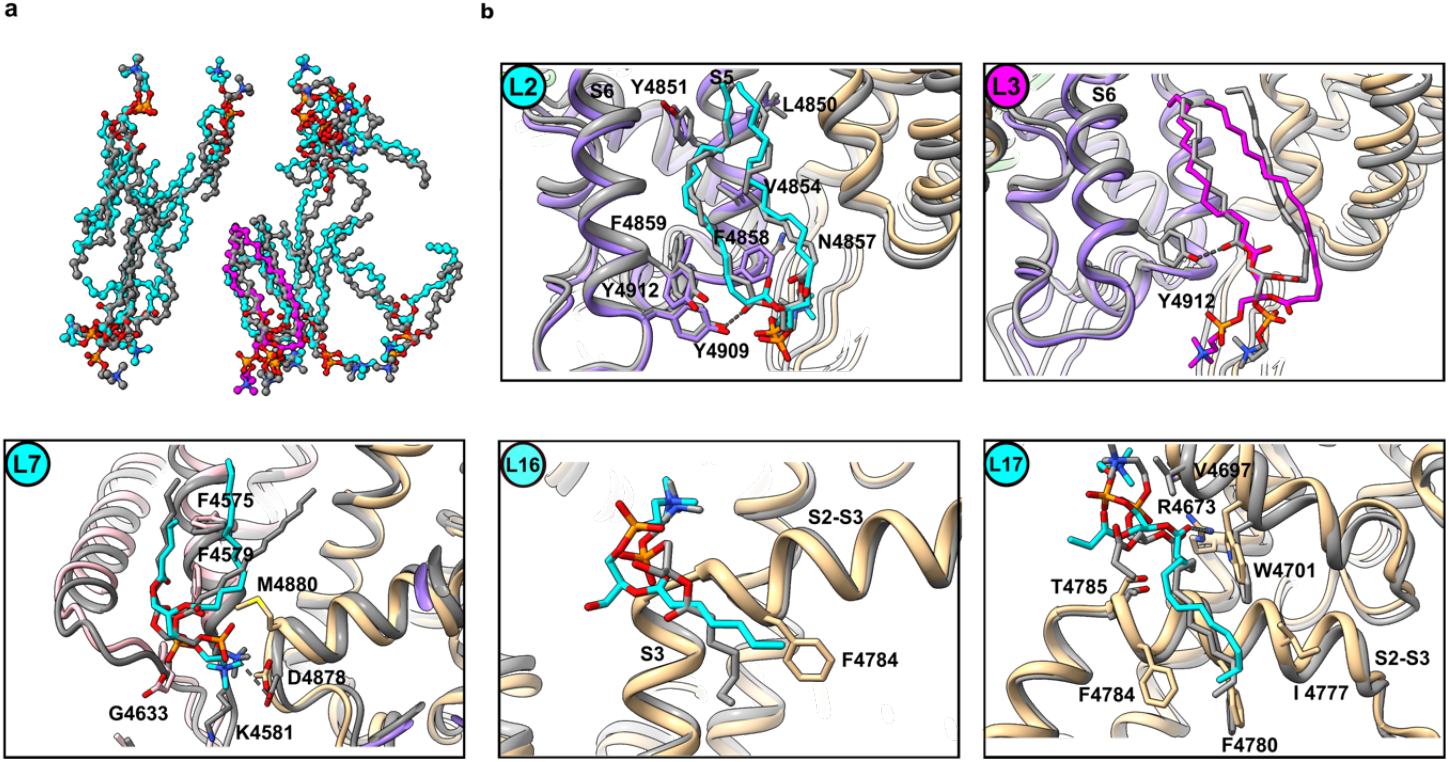
Structural differences in lipid positions between primed and open states. **a**, Comparison of lipid models between the DT-0.05-open and DT-0.05-primed states. **b**, Detailed view of lipid rearrangements. The structures were aligned on the pore region (residues 4820-5037). The open state is shown in grey (For clarity, only lipids conserved in the primed state are shown), and the primed state is colored as in **Figure 3**.

### Open probability correlates with dimensions of the bilayer-mimicking belt

An additional insight into the properties of lipid bilayer mimetics was obtained from the analysis of the shape of the belt observed in the cryo-EM maps around membrane domain of RyR1. This hydrophobic belt originates from the lipid bilayer mimetics and reflects its geometry. We compared the sizes of the nanodiscs, micelles, and liposomes for nine density maps (**Figure 5 a**). For this analysis, the pixel size of the maps was calibrated, the maps were low pass filtered to resolution of 8 Å and the density levels were set to enclose identical volumes of the cytoplasmic domains of the RyR1 reconstructions.

**Figure 5.**
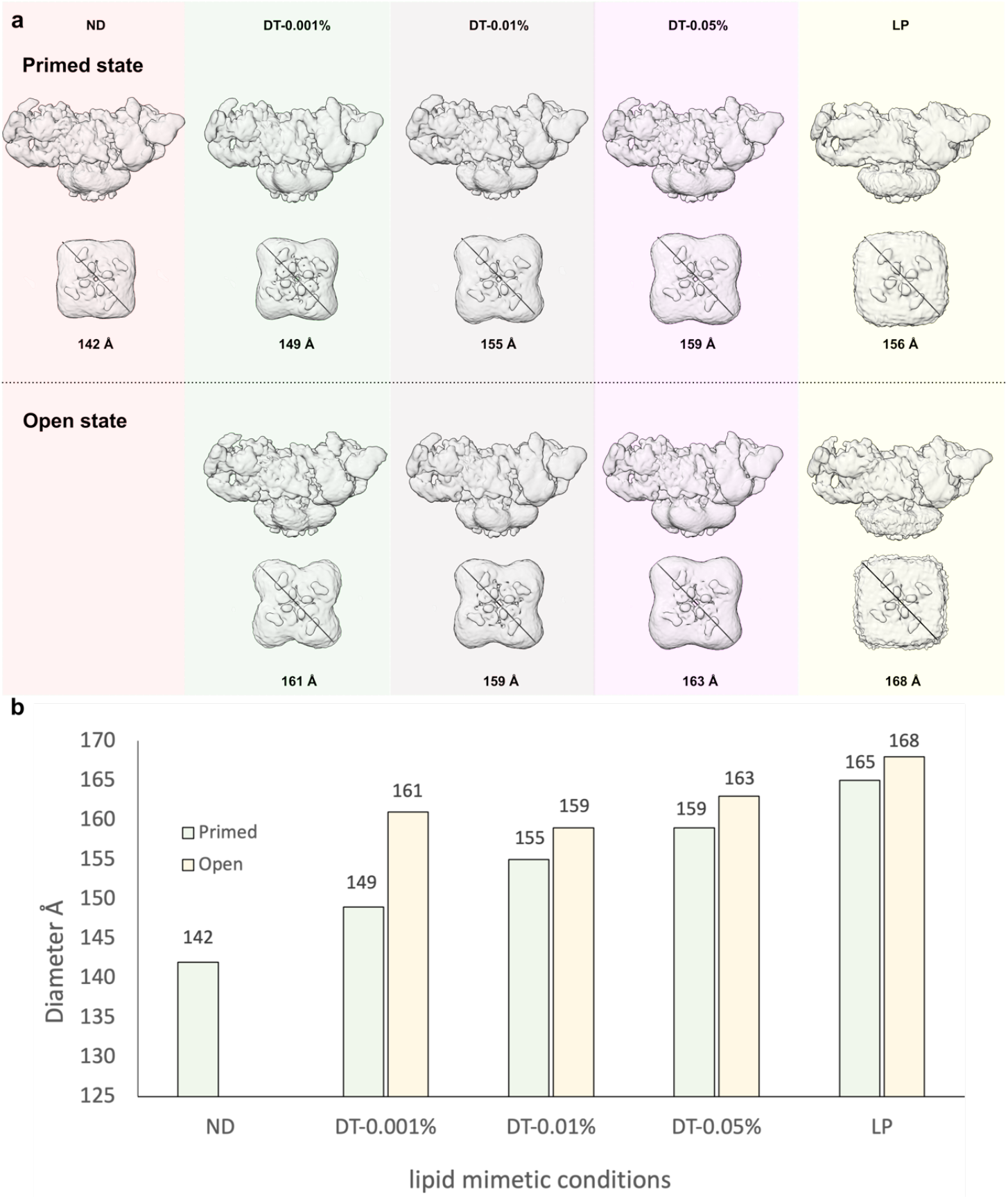
Maps of hydrophobic belts for RyR1 reconstituted in various lipid bilayer mimetics. **a**, Density maps of RyR1 reconstituted in various lipid bilayer mimetics. From left to right: ND, DT-0.001, DT-0.01, DT-0.05, and LP. The first two rows display side and bottom views of the primed state maps, while the third and fourth rows present side and bottom views of the open state maps. The diameter of the bilayer-mimicking belt for maps shown in **a**.

The nanodisc belt had the smallest diameter of 142 Å (average value of three measurements, other diameters are calculated in the same way). For detergent-solubilized RyR1, the micelle sizes increased with lipid concentration from ∼149 Å at 0.001% to 159 Å at 0.05% POPC, respectively. The consistent density of liposome-reconstituted RyR1 had a diameter of ∼156 Å **(Figure 5)**.

In the open state, the size of the micelle increased by 5.8+/-4.2 Å. This is consistent with outward rotation of the voltage-sensor-like lobes with amplitude of ∼5 Å (diameter increase of the modeled region from 131 to 136 Å). As lipid concentration increased, the grooves between S1-S4 lobes were less pronounced and overall shape of the micelle became rounder (**Figure 5 a**). This is consistent with binding of more lipids in the grooves as was described in the previous section. Taken together, these data indicate that the fraction of open RyR1 particles correlates with the size of the belt mimicking lipids bilayer. The LP datasets are likely less representative than lipid bilayer mimetics because lipid bilayer there is continuous, and the size of the belt reflects consistency of the liposome curvature rather than their physical dimensions.

## Discussion

Gating of ion channel is a stochastic process, thus, even in the closed state RyR1 shows rare transient spikes of conductivity ^8,53^. Channel activators modulate frequency of conductivity spikes and their length ^54^. Cryo-EM captures statistically averaged conformations which are expected to reflect the time-averages of open and closed channel states. There is an agreement between electrophysiology and cryo-EM regarding the existence of 2 principal channel conductivity states: closed pore that allows no current flow and open pore conformation in which hydrophobic gate made by side chains of I4937 is open to a well-defined diameter that facilitates ion flow with a certain resistance. Where cryo-EM data consistently disagree with electrophysiology for RyR1 is the channel open probability which consistently produced lower estimates from cryo-EM data than observed by single channel recordings.

Here we show that when RyR1 is solubilized in CHAPS, its open conformation can be modulated by lipids in a broad range. When lipid concentration is varied 50-fold between 0.001 and 0.05%, open probability of RyR1 increases from 16 to 84%. In this range the molar ratio between CHAPS molecules and POPC changes from 250:1 to 5:1 meaning that the starting point represents nearly pure CHAPS micelles whereas at high lipid concentration, about 20% of the molecules in a detergent micelle are POPC. At high lipid concentration, the open probability of RyR1 calculated from cryo-EM closely matches the open probability of RyR1 observed in the electrophysiological experiments with the same channel activators Po ∼0.96 ^8^ suggesting that 5:1 CHAPS:POPC micelles provide a native-like functional environment for RyR1. It is worth noting that addition of lipids to detergent solution reduces the contrast of protein particles in the electron micrographs and therefore usually avoided during preparations of cryo-EM grids, but as we show here, it is desirable in specific cases like the studies of the open conformation of RyR1.

The properties of detergent micelles are very different from those of lipid bilayer. Detergent molecules are less hydrophobic, have different geometry, charge distribution and shape. Hydration of the detergent micelles is different from that of lipid bilayer. Therefore, it is not surprising that the free energies of the conformations associated with channel gating are different in lipid bilayer and in detergent micelles.

But what is the mechanism by which lipids stabilize the open conformation of RyR1 in detergent micelles? Cryo-EM structures suggest that addition of lipids alters structure of the primed state only slightly but the number of ordered lipids increases significantly. At high concentration of lipids, grooves between the voltage-sensor-like domains of RyR1 are filled with two layers of nearly complete bilayer-like lipids structure. These are also the regions where the largest conformational changes take place on the cytoplasmic surface of the membrane domain around trans-membrane helices S5 and S6. Hence, these lipids provide very local native-like bilayer environment around the gate and may improve hydrophobicity of the micelle in these regions as well as create native-like local electrostatic potential. The size of the micelles also increases providing more hydrophobic environment to the trans-membrane domain as a whole which is also consistent with rendering the micelle more lipid bilayer like. Stabilization of the closed state at low lipid concentration in the CHAPS micelles is also consistent with the observation that at low lipid content the structures of the gate region are somewhat tighter in the primed and closed states than at high lipid concentration.

Our study compares 5 RyR1 structures all bound to a nanobody used for protein purification ^45^. Even though the nanobody was bound to Repeat12 domain at the periphery of the 270 Å wide cytoplasmic crown of RyR1, we cannot exclude a systematic effect of the nanobody on the open fraction of the channels, particularly because dantrolene and ARM210 bind to the overlapping sites ^43,55^. This, however, does not undermine our conclusions because all conditions apart from lipid mimetics were identical for all the compared structures.

It must be also noted that our study relied of miniaturized purification protocol that does not involve commonly used spin concentrators. This ensured accurate control of lipid concentration in the cryo-EM samples. Protein concentrating using porous-membrane-based spin concentrators, might lead to variability of the final detergent and lipid concentration ^56^ and consequently to the variability of RyR1 open probability between preparations. The lipid and detergent concentrating effect may explain the inconsistent between our results (Po= 16% at 0.001% POPC) and the study reported by de Georges at.al. ^8^ (Po=50%) that used spin concentrator during the last step of the protein preparation.

Intriguingly, reconstituted into lipid nanodiscs, RyR1 was expected to be in lipid bilayer like environment ^57,58^, however, the channel remained fully closed despite various attempts to open it. RyR1 was reconstituted into lipid nanodiscs using membrane scaffold protein MSP1E3D1 and POPC. To reduce aggregation of the reconstituted RyR1, we added 0.06% fluorinated octyl maltoside (fOM) ^45^. To open the channel, we attempted to remove fOM and to change the lipid composition of the nanodiscs (SoyPC, and a 1:1 mix of POPC and DLPC mixture ^59^ were tried), but the channel remained closed. We know however, that ryanodine does stabilize RyR1-ND in fully open state ^24^ suggesting that RyR1-ND can be opened but requires a stronger agonist. Consistently, RyR1-ND had the smallest hydrophobic belt among the studied lipid mimetics (**Figure 5**). The estimated size of the nanodisc of 142 Å is consistent with the expected diameter of MSP1E3D1-based nanodisc (120 Å to 160 Å ^60–62^) and its shape suggests that the grooves between voltage-sensor-like domains are filled with lipids as would be expected for the nanodisc. Despite this, the ordered lipids were not well-resolved in the RyR1-ND cryo-EM map even though it res reconstructed to a higher resolution. We can hypothesize that in the nanodisc, the lipid belt constrained by MSP1E3D1 is too thin to mimic bulk of a lipid bilayer and lipid-MSP interaction propagates as a disorder in the grooves between voltage-sensor-like domains.

Occupancy of the open channel conformation of RyR1 reconstituted into lipid vesicles was in the range 37% and 62%, which is also lower than expected from single channel recordings and as previously discussed ^45^ may depend on the lipid composition of liposomes and residual amounts of CHAPS in the reconstituted sample. Additionally, the high lipid bilayer curvature may contribute to different lipid pressure on the luminal and cytoplasmic sides of the membrane.

In summary, this report shows that open probability of RyR1 is highly sensitive to the lipid bilayer mimetics and must be thoroughly controlled during sample preparation for cryo-EM studies that investigate fine regulation of RyR1 gating by modulators.

## Methods

### Rabbit skeletal SR membranes isolation

The method used for SR membrane isolation here are adapted from a previously described protocol ^63^. All steps were conducted on ice or at 4°C. Around 120 g of muscle tissue from a male New Zealand White rabbit of 8 weeks old was cut into 1 cm^3^ and homogenized in ice cold buffer (20 mM Tris-maleate pH 7.0, 100 mM NaCl, 0.3 M sucrose, 1 mM EGTA, 2 mM DTT, and a cocktail of inhibitors: 4 mM leupeptine, 1 mM benzamidine, 0.1 mM aprotinin, 1 mM pepstatin, 2 mM calpain I inhibitor and 1 mM phenylmethylsulphonyl fluoride (PMSF)) to a total volume of 1.6 l using a Waring blender. The homogenization was performed in bursts with duration of 30 s each until the homogenized suspension appeared pink and smooth. Then the suspension was centrifuged 30 min at 7,000 rpm, in Beckman *JA-10* rotor. The supernatant was filtered through two layers of cheesecloth, 0.5 M of KCl was added and stirred for at least 10 min. The extract was centrifuged at 40,000 rpm, 30 min in Beckman 45Ti rotor. The pellet was resuspended in buffer containing 20 mM MOPS, pH 7.4, 1 mM EGTA, 2 mM DTT, 0.6 M KCl, 0.3 M sucrose and cocktail of inhibitors to a final volume of 80-100 ml using a glass homogenizer and centrifuged at 40,000 rpm for 30 min in Beckman 45Ti rotor. The pellets were homogenized in the resuspension buffer to a concentration of around 10 mg/ml, frozen in liquid nitrogen and stored at -80°C. Protein concentration was determined by Pierce 660 nm Protein assay (Thermo Scientific).

### Preparation of periplasmic extract containing Nbs

The nanobody repertoire was cloned into the phage-display vector pMESy4, facilitating the expression of Nbs with a 6xHis-tag. Overexpression of Nbs occurred overnight following induction with 1 mM Isopropyl β-D-1-thiogalactopyranoside (IPTG) in Escherichia coli WK6 strain, initiated once the cell density reached OD 0.8 at 600 nm. Cells were harvested by centrifugation at 5,000 rpm for 15 min using a Beckman JA-10 rotor. The cell pellet was lysed by adding 4 mL of lysis buffer (50 mM Tris pH 7.5, 150 mM NaCl, 1 mM EDTA, 0.1 mg/ml lysozyme, 20% sucrose, 50 µg/ml DNase, and 1 mM PMSF) per 1 g of pellet, followed by a 30 min incubation at 4°C. Periplasmic extract was obtained by centrifugation at 10,000 rpm for 15 min using a Beckman JA-20 rotor. MgCl^2^ (5 mM) was added to the extract followed by filtration through 0.45 µm Millipore filter and supplementation with 20% sucrose before storage at -80°C in aliquots.

### Nanobody-based affinity RyR1 purification

Every step of the purification was performed at 4°C or on ice. 0.5 ml of SR membranes (10 mg/ml) was solubilized by 375 µl solubilization buffer (20 mM MOPS pH7.4, 1 M NaCl, 10% sucrose, 2 mM TECP, 4% CHAPS 0.8% POPC, and a cocktail of protease inhibitors) for at least 30 min following 30 min ultracentrifugation at 40, 000 rpm in Beckman 50.4 Ti rotor. Then, the supernatant was incubated with nanobody loaded magnetic beads for 30 min and washed with SBL buffer (similar to solubilization buffer except the NaCl concentration is 0.7 M, CHAPS and POPC, concentration are 0.8% and 0.2%, and add 200 µM EGTA). Nanobody-loaded magnetic beads (HisPur™ Ni-NTA Magnetic Beads, Thermo Fisher Scientific) were prepared by incubating 135 µl washed magnetic beads with 135 µl Nb9657-His periplasmic extract for 15 min. Magnetic beads were equilibrated and washed with SBL buffer by adding 270 µl SBL buffer. First elution was done by incubating 65 µl elution buffer1 (SBL buffer with 500 mM Imidazole) with magnetic beads for 5 min. At the second purification step, 65µl of eluate was diluted with 600 µl SBL buffer and the mix incubated for 30 min with another 135 µl of Nb9657 loaded magnetic beads. After incubation, the beads were washed three times with SBL buffer.

Second elution was performed by incubating the beads with 60 µl of elution buffer 2 (SBL buffer with 2 mM ATP, 5 mM caffeine and 50 µM free Ca^2+^ and 500 mM Imidazole) followed by separating the beads from the eluate using a magnetic stand. The elution 2 was then applied to the homemade spin column equilibrated in desalting buffer 1 (elution buffer 2 without Imidazole) and centrifuged for 1 min at 1000 xg to remove Imidazole. After first desalting step, 50 µl of eluate was incubate with 60 µg of Mb9657 for at least 1h. The desalting was repeated using desalting buffer2 (similar desalting buffer1, the NaCl concentration was 0.2 M with either 0.2% CHPAS and 0.05% POPC or 0.2% CHAPS and 0.01% POPC).

### Preparation of graphene oxide grids

Home-made graphene oxide grids were utilized for sample preparation. A stock solution was created by diluting a 2 mg/ml graphene oxide solution (Sigma) with Milli-Q water to achieve a concentration of 0.6 mg/ml. Prior to each use, the stock solution underwent centrifugation for 3 minutes at 3000 x g, followed by a 5-fold dilution with Milli-Q water to yield a final concentration of 0.12 mg/ml. Quantifoil R2/1 Cu300 holey carbon grids were subjected to glow discharge in the ELMO glow discharge system (Corduan Technologies) from the carbon side for 1 minute at 5 mA and 0.28 mbar. Subsequently, 3µl of the 0.12 mg/ml graphene oxide solution was applied to the carbon side of the grids and incubated for a minimum of 1 minute. The grids were then gently blotted with Whatman filter paper No. 1, followed by a wash with 10 µl of distilled water and immediate blotting with Whatman No. 1 filter paper. Finally, the grids were air-dried for at least 30 minutes at room temperature.

### Preparation of cryo-EM grids

Home-made graphene oxide grids were prepared as described elsewhere ^45^. The cryo-EM samples were prepared using a CP3 cryoplunge system (Gatan). A 4 µl aliquot of the protein solution was applied to the carbon side, while 1 µl of desalting buffer without lipids was added to the copper side of the grid. Blotting from both sides was performed for 1.2 seconds using Whatman glass microfiber filter paper GF/A at 91% relative humidity. The grids were plunge-frozen in liquid ethane at -175°C and then stored in liquid nitrogen.

### Data collection

Cryo-EM images were acquired using a JEOL CryoARM300 microscope, with in-column Ω energy filter and a cold field emission gun (cFEG) operating at 300 kV^64^. Images were collected on K3 detector (Gatan). Movies comprising 60 frames at a nominal magnification of 60,000x were automatically captured using SerialEM 3.0.8 ^65^. The energy filter slit was set to a width of 20 eV. Each frame had an exposure time of 2.796 seconds, with an average dose per frame of 1 electron per square angstrom (1 e-/Å^2^), and defocus values ranging from -1.5 to -2.5 µm. The pixel sizes ranged from 0.73 Å to 0.76 Å. A cross-shaped pattern was used to collect 25 images per single stage position (5 per hole), with three holes along each axis ^66^.

### Data processing

For all datasets, motion correction was performed using MotionCor2 ^67^. Contrast Transfer Function (CTF) parameters were estimated using CTFFIND-4.1^68^. Subsequently, all images were imported into cryoSPARC (versions 4.6.2) ^69^ for further processing. Particle picking was conducted with template picker using 2D templates created from already processed datasets. Following particle picking, all particles were extracted into boxes of 168 pixels, with pixel size of 2.92 Å. Multiple rounds of 2D classification were then performed to select particles for subsequent processing (**Supplementary Figures 1 and 2**).

For the ND-0.01 dataset, 128,894 particles were selected after 2D classification to generate an initial model with C1 symmetry. A total of 127,957 particles were re-extracted into 336-pixel boxes with a pixel size of 1.46 Å. These particles were refined into a consensus 3D map with C4 symmetry, achieving a resolution of 3.6 Å. The particles were imported into Relion v4.0.1 for polishing and 3D classification.

During the 3D classification, particles were divided into six classes using a mask focused on the transmembrane region (**Supplementary Figure 1**) to separate the open and primed states. Classes 1, 2, 4, and 5 represented the open conformation, comprising 80,709 particles. Class 3, containing 14,320 particles, corresponded to the primed conformation, while particles contributing to class 6, with poor density, were discarded. The open and primed state particles were imported back into cryoSPARC v4.6.2 for refinement using the NU-Refinement procedure. The consensus maps of the open and primed states were reconstructed to resolutions of 3.6 Å and 3.3 Å, respectively.

To improve the density of flexible domains, four local refinements were performed using local masks (**Supplementary Figure 1**). Mask 1 covered TaF, TM, and CTD domains, mask 2-JSolA and CSol, mask 3-FKBP and TTD, and mask 4-BSol.

For the open state, local refinement with mask 1 was applied to the 80,709 particles with C4 symmetry. Masks 2, 3, and 4 were used for local refinement with C1 symmetry on 322,836 particles generated through C4 symmetry expansion of the 80,709 particles. The resulting local maps had resolutions of 3.1 Å (mask 1), 3.2 Å (mask 2), 3.1 Å (mask 3), and 3.9 Å (mask 4).

For the primed state, local refinement with mask 1 was performed on 39,983 particles with C4 symmetry. Masks 2, 3, and 4 were used for local refinement with C1 symmetry on 159,932 particles obtained after C4 symmetry expansion of the 39,983 particles. The resulting local maps had resolutions of 3.4 Å (mask 1), 3.3 Å (mask 2), 3.1 Å (mask 3), and 3.8 Å (mask 4).

The resulting maps were combined in Chimera v1.15 to generate a composite map. Pixel size calibration and model building were performed on the composite map.

The DT-0.05 dataset was processed similarly (**Supplementary Figure 2**). A total of 335,432 particles were selected after 2D classification and used to generate an ab initio model with C1 symmetry (**Supplementary Figure 2**). The particles were re-extracted into 336-pixel boxes at a pixel size of 1.51 Å and refined into a consensus 3D map with C4 symmetry.

All particles were imported into Relion v4.0.1 for polishing and 3D classification. Six classes were defined for classification: classes 1, 4, and 5 represented the open conformation and comprised 279,776 particles. Class 6 corresponded to the primed conformation, comprising 53,158 particles. Classes 2 and 3 were discarded due to poor density. The consensus map of the open state was reconstructed to a resolution of 3.4 Å, with local maps achieving resolutions of 3.3 Å for masks 1, 2, and 3, and 3.6 Å for mask 4. The consensus map of the primed state was reconstructed to a resolution of 3.6 Å with local maps achieving resolutions of 3.3 Å for mask 1, 3.5 Å for mask 2, 3.3 Å for mask 3, and 4.4 Å for mask 4.

### Model building

Initial model of open state PDB: 8RRW, and 8RRX for primed state were used. The initial models were rigid body fit to each map in Chimera v 1.15 ^70^ then manually rebuilt in Coot 0.98 ^71,72^ followed by real-space refinement in Phenix 1.20.1 using default parameters. All models were validated with MolProbity ^73^. The figures were prepared using ChimeraX-1.7.1 ^70^.

## Acknowledgements

We thank Annelore Stroobants for technical support, Dr. Marcus Fislage for assistance with cryo-EM data collections. This work was funded by grants from FWO (grant Nos. G.0266.15N, G0H5916N, G054617N to R.G.E.) and the European Research Council (Grant No. 726436 to R.G.E).

## Notes

### Competing Interest Statement

The authors have declared no competing interest.

