## Supplementary Information for "Lipids modulate open probability of RyR1 under cryo-EM conditions"

[*] Corresponding author

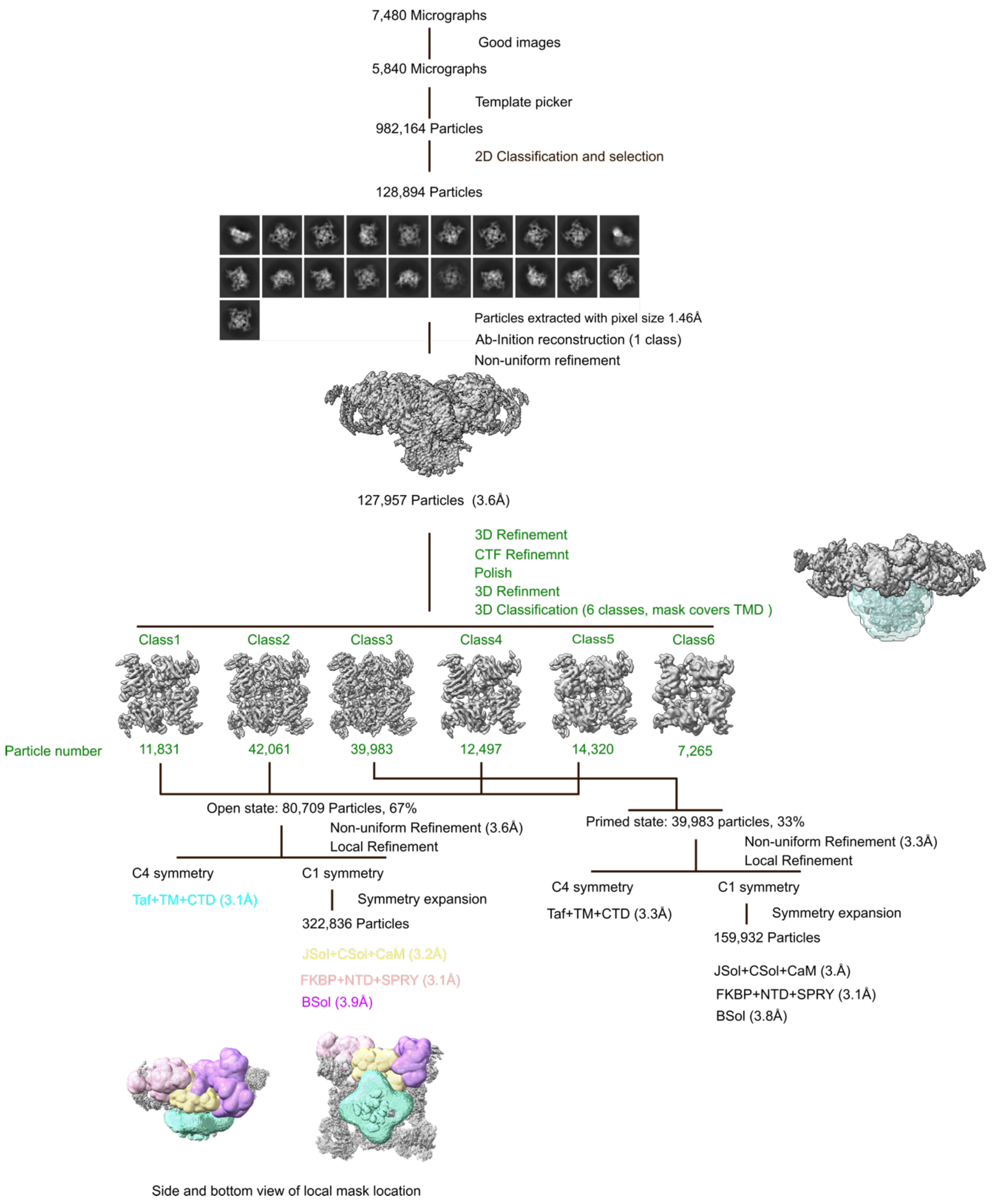


**Supplementary Figure1 | Data processing of 0.01% POPC dataset.** Green color indicates steps performed in Relion, other steps were performed in CryoSPARC. The masks used for 3D classification are indicated next to the 3D classification step. The masks used for local refinement are shown at the bottom.

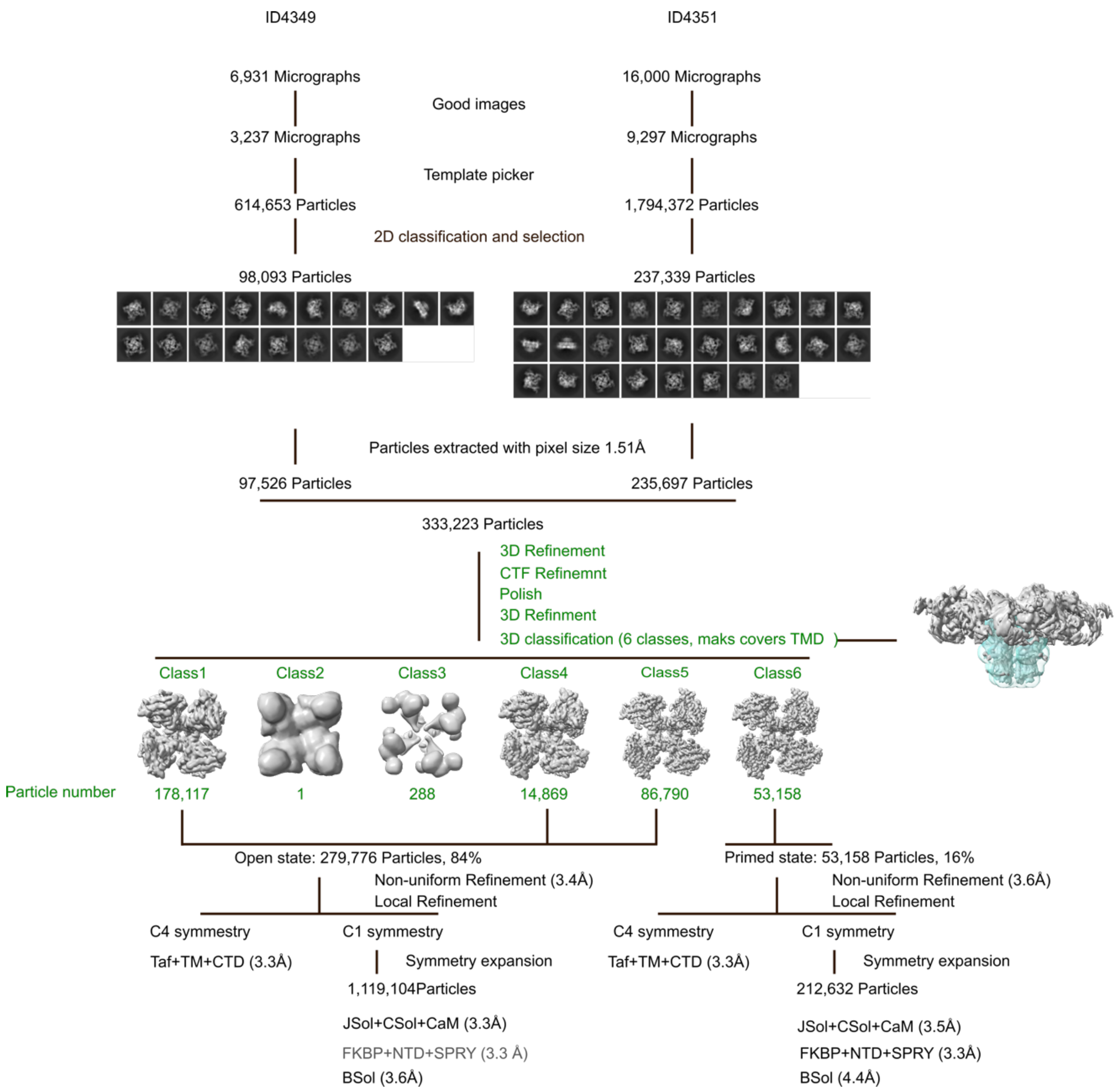


**Supplementary Figure 2 | Data processing of 0.05% POPC dataset**.

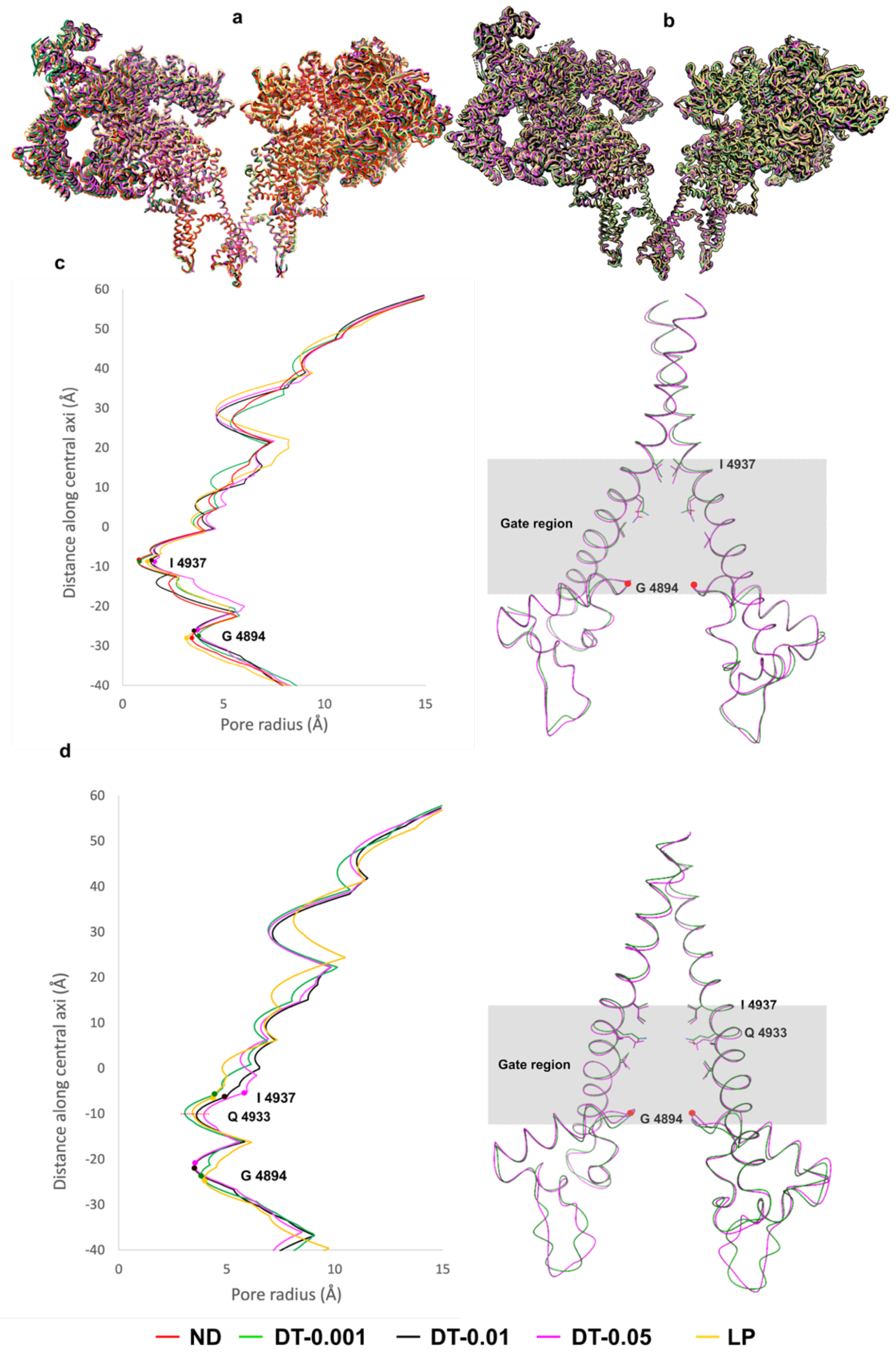


**Supplementary Figure 3 | Structure of RyR1 pore in different lipid mimicking environments. a,b,** Comparison of structures in primed state and open state. The structures were aligned on the pore region (residue 4820-5037). Model colors are indicated by a legend at the bottom of the image. **c, d,** The pore radius and pore regions structure in the primed and open states, respectively. On the left, the pore radius, calculated using HOLE, is shown. Gate residue I4937 and selectivity filter residue G4984 are marked by dots. On the right, the structures of the pore regions are shown. For clarity, only DT-0.001 and DT-0.05, which show the greatest differences, are displayed.

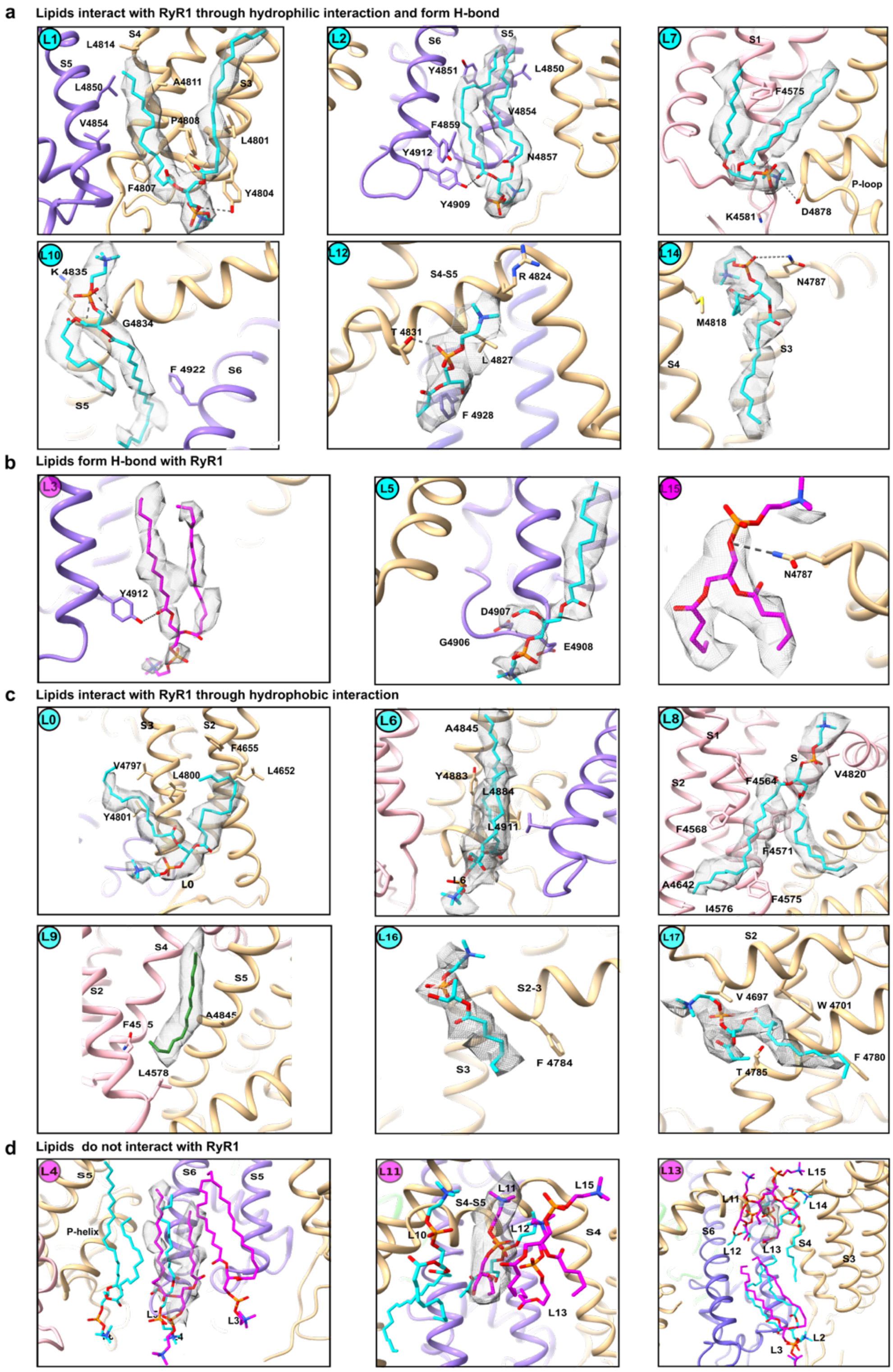


**Supplementary Figure 4 | Interaction of individual lipids with RyR1.** The RyR1 is shown in cartoon representation. Protein 4 subunits are colored brown, pink, lime, and purple. Lipids are shown as sticks and colored the same way as in **Figure 4**; the lipid number is labeled in the top left corner. The density of the analyzed lipid is shown as a gray mesh. **a, b, c**, Show lipids interacting with RyR1 through both hydrophobic contacts and hydrogen bonds, only hydrogen bonds, and only hydrophobic interactions, respectively. **d**, Lipids without direct interactions with RyR1.

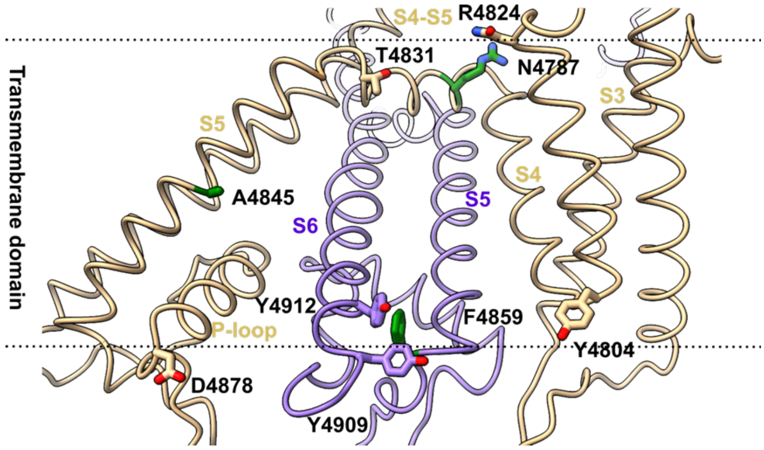


**Supplementary Figure 5 | Residues involved in polar RyR1-lipid interactions**. For clarity, only two neighboring protomers are shown; the polar side chains making putative interactions with lipids are shown as sticks. Residues mutated in diseases are shown in green.

### Supplementary Table 1 | Statistics of cryo-EM data and models.

| **Sample** | **0.05% lipids** | | **0.01% lipids** | |
|---|---|---|---|---|
| State | open | primed | open | primed |
| Ligands | 50µM free $Ca^{2+}$, 2mM ATP, 5mM Caffeine | | | |
| PDB ID | 9HEO | 9HEP | 9HEQ | 9HER |
| EMBD ID | EMD-52085 | EMD-52088 | EMD-52091 | EMD-52094 |
| | **Data collection** | | | |
| Microscope | JOEL CRYOARM300 | | | |
| Detector | Gatan K3 | | | |
| Voltage(kV) | 300 | | | |
| Magnification | 60,000 | | | |
| Exposure time (S) | 2,796 | | | |
| Electro dose (e -/Å $^2$) | 60 | | | |
| Number of frames | 60 | | | |
| Defocus range (µm) | 1.5-2.5 | | | |
| Pixel size (Å) | 0.76 | | | |
| Symmetry | C4 | | | |
| Collected images | 22,931 | | 7480 | |
| Used images | 12,534 | | 5840 | |
| Final particles (N) | 279,776 | 53,158 | 80,709 | 39,983 |
| Global map resolution (Å) | 3.4 | 3.6 | 3.6 | 3.3 |
| Local maps resolution range (Å) | 3.3-3.6 | 3.3-4.4 | 3.1-3.9 | 3-3.8 |
| **Model refinement** | | | | |
| Refinement package | Phenix v 1.20.1 | | | |
| Initial model used | | | | |
| Model resolution [Å$^2$], FSC=0.5 | 3.4 | 3.6 | 3.6 | 3.4 |
| **Model composition** | | | | |
| Non-hydrogen protein atoms | 146,795 | 145,746 | 145,720 | 145,068 |
| Protein residues | 18,244 | 18,224 | 18,204 | 18,204 |
| Ligands | 88 | 64 | 64 | 52 |
| **B-factors mean [Å$^2$]** | | | | |
| Protein | 121.29 | 113.85 | 107.23 | 145.84 |
| Ligand | 110.39 | 104.33 | 192.67 | 168.20 |
| **R.M.S deviations** | | | | |
| Bond lengths (Å) | 0.004 | 0.003 | 0.003 | 0.003 |
| Bond angles (°) | 0.694 | 0.725 | 0.758 | 0.703 |
| **Validation** | | | | |
| Molprobity score | 1.68 | 1.67 | 1.65 | 1.6 |
| Clashscore | 9.54 | 9.80 | 10.76 | 8.35 |
| Poor rotamers (%) | 0.52 | 0.39 | 0.62 | 0.31 |
| **Ramachandran plot** | | | | |
| Favored (%) | 96.99 | 97.17 | 97.54 | 97.23 |
| Allowed (%) | 2.94 | 2.77 | 2.44 | 2.75 |
| Disallowed (%) | 0.07 | 0.07 | 0.02 | 0.02 |

### Supplementary Table 2 | Open fractions of rabbit RyR1 in different lipid mimetics

| **Sample** | **ND** | **DT-0.001%** | | **DT-0.01%** | | **DT-0.05%** | | **LP** | |
|---|---|---|---|---|---|---|---|---|---|
| Ligands | 50µM free $Ca^{2+}$, 2mM ATP, 5mM Caffeine | | | | | | | | |
| state | primed | open | primed | open | primed | open | primed | open | primed |
| Resolution (Å) | 3.1 | 4.2 | 3.3 | 3.6 | 3.3 | 3.4 | 3.6 | 4.6 | 4.7 |
| Final particles (N) | 166,010 | 29,246 | 145,830 | 80,709 | 39,983 | 279,776 | 53,158 | 26,815 | 16,530 |
| Open particle percentage (%) | 0 | 16 | | 67 | | 84 | | 62 | |

**Supplementary Table 3 | Lipid Statistics in Structures**. The cell color indicates the layer where the lipid is located: cyan for the first layer, and magenta for the second layer. Text color indicates the type of interaction with RyR1 (based on the DT-0.05% structure): red for hydrogen bonds, blue for both hydrogen bonds and hydrophobic interactions, yellow for hydrophobic interactions, and black for no interaction. Blank cells indicate that the lipid does not exist in that specific structure.

| Structure | Lipid number | | | | | | | | | | | | | | | | | |
|---|---|---|---|---|---|---|---|---|---|---|---|---|---|---|---|---|---|---|
| 0.05 open | L0 | L1 | L2 | L3 | L4 | L5 | L6 | L7 | L8 | L9 | L10 | L11 | L12 | L13 | L14 | L15 | L16 | L17 |
| 0.05 primed | L0 | L1 | L2 | L3 | | | L6 | L7 | L8 | L9 | L10 | | | | L14 | | L16 | L17 |
| 0.01 open | L0 | L1 | L2 | L3 | | | L6 | L7 | L8 | L9 | L10 | L11 | | | L14 | | | L17 |
| 0.01 primed | L0 | L1 | L2 | | | | L6 | L7 | | L9 | L10 | | | | L14 | | | L17 |
| 0.001 primed | L0 | L1 | L2 | L3 | | L5 | L6 | L7 | L8 | L9 | L10 | L11 | | L13 | L14 | | L16 | L17 |
| ND-primed | L0 | L1 | L2 | L3 | | | L6 | L7 | L8 | L9 | L10 | | L12 | | L14 | | | L7 |

**Supplementary Table 4 | Correlation coefficient statistics of lipids across different structures.** The values were estimated in Phenix Comprehensive validation (cryo-EM). The cross-correlation coefficient was calculated only for lipids that were resolved in the corresponding map

| Structure | DT-0.05-open | DT-0.05-primed | DT-0.01-open | DT-0.01-primed | DT-0.001-primed | ND |
|---|---|---|---|---|---|---|
| CC | 0.66 | 0.56 | 0.62 | 0.57 | 0.42 | 0.39 |

### Supplementary Table 5 | Key resource data

| Deposited Data | Source | Identifier |
|---|---|---|
| RyR1-ND | Li et al, 2024 | PDB: 8RRX<br>EMDB: 19468 |
| RyR1-0.001-primed | Li et al, 2024 | PDB: 8RSO<br>EMDB: 19472 |
| RyR1-0.001-open | Li et al, 2024 | PDB: 8RRW<br>EMDB: 19467 |
| RyR1-0.01-primed | This study | PDB:9HER<br>EMDB:52094 |
| RyR1-0.01-open | This study | PDB:9HEQ<br>EMDB:52091 |
| RyR1-0.05-primed | This study | PDB:9HEP<br>EMDB:52088 |
| RyR1-0.05-open | This study | PDB:9HEO<br>EMDB:52085 |
| RyR1-LP-primed | Li et al, 2024 | PDB:8RRU<br>EMDB: 19465 |

**Supplementary Video 1 | Conformational change in RyR1 and modelled lipids between primed and open states.**
